# Father-child dyads exhibit unique inter-subject synchronisation during co-viewing of animation video stimuli

**DOI:** 10.1101/2020.10.30.361592

**Authors:** Atiqah Azhari, Andrea Bizzego, Gianluca Esposito

**Affiliations:** Psychology Program, School of Social Sciences, Nanyang Technological University, Singapore; Department of Psychology and Cognitive Science, University of Trento, Italy; LKC School of Medicine, Nanyang Technological University, Singapore

**Keywords:** synchrony, inter-subject synchronisation, parent-child, father-child, NIRS, prefrontal cortex, narrative scene, video stimuli, co-viewing

## Abstract

Inter-subject synchronisation reflects the entrainment of two individuals to each other’s brain signals during passive joint tasks. Within the parent-child dyad, the temporal coordination of signals indicates an attunement to each other’s emotional states. Despite the ubiquity with which parents and their children watch screen media together, no study has investigated intersubject synchronisation in father-child dyads during such a co-viewing activity. The present study examined whether father-child dyads would exhibit unique inter-subject synchronisation during co-viewing of narrative visual scenes that is unique to the dyad and hence would not be observed in control dyads (i.e. randomly paired signals). Hyperscanning fNIRS was used to record the prefrontal cortex (PFC) signals of 29 fathers and their preschool-aged children (11 girls, 18 boys) as each pair engaged in a typical activity of watching children’s shows together. Three 1-min video clips from “Brave”, “Peppa Pig” and “The Incredibles” were presented to each dyad and children’s ratings of video positivity and familiarity were obtained afterwards. PFC activity was analysed according to four clusters: medial left, medial right, frontal left and frontal right clusters. Results from synchrony analyses demonstrated that true father-child dyads showed significantly greater inter-subject synchronisation than control dyads, especially in the medial left cluster during the emotionally arousing conflict scene. Dyads with older fathers displayed less synchrony and older fathers, compared to younger ones, were also found to exhibit greater activity in the frontal right cluster. These findings point to a unique inter-subject synchronisation that exists in father-child relationships during shared co-viewing of narrative scenes which could be potentially modulated by parental age.

## INTRODUCTION

At the turn of the millennium, the predominant single-brain paradigm in social neuroscience which focused on the individual brain was complemented with a burgeoning dual-brain approach. Investigating two brains in tandem enabled unparalleled inquiry into how similarly the world is being processed by different people [1]. Captivated by this possibility, [2] examined how visual stimuli in a naturalistic environment, in the form of a narrative movie, might elicit similar brain signals across individuals. Their seminal paper discovered the phenomenon of inter-subject synchronisation in which the brain signals of two or more persons are temporally coordinated during passive joint activities like co-viewing. Synchronous activation patterns were not only detected in visual and auditory networks well-established to be implicated in sensory mechanisms, but were also evident in other higher-order cortical areas. These observations led to the conclusion that the narrative scene functioned as an environmental stimulus which entrained the brain signals of individuals to elicit inter-subject synchronisation. A considerable number of studies have since employed a similar paradigm of presenting participants with narrative scenes or audios, which lent further support to the occurrence of intersubject synchronisation [2]–[7].

Several mechanisms by which intersubject synchronisation occurs have been posited. Using an analogy of a “wireless transmission”,[5] described how two systems, akin to two persons, correspond with each other via a shared signal which manifests through mutually seeing, hearing or feeling each other’s responses towards the same environmental stimulus. Receiving input from another person who inhabits a comparable human physical form is thought to activate vicarious brain activations which underpin similar brain responses [5], [8]. Buttressing this explanation, [9] proposed that the exposure to narrative movies trigger a common script that is shared across different people. For instance, viewing a conflict scene activates a similar set of antecedents and expectations associated with relational discord that are arguably distinct from watching an amicable scene. Hence, the shared script that individuals process narrative information with, along with the resemblance that individuals have to each other which activates similar vicarious brain responses, are postulated to drive inter-subject synchronisation.

Compared to other areas of the brain, the prefrontal cortex showed greater variability in the extent to which inter-subject synchronisation was observed [2]. This disparity was thought to be rooted in interindividual differences in processing the same narrative scene. However, it remains to be seen whether individuals within a close attachment relationship, such as that of a parent-child dyad, would show significantly greater synchrony compared to that observed between two arbitrary individuals.

The maturing parent-child relationship presents dyads with innumerable opportunities for partners to develop biobehavioural synchrony [10], [11]. Whereas inter-subject synchronisation measures the alignment of brain signals during a passive task, biobehavioural synchrony is a construct that reflects the reciprocal matching of signals at both the behavioural and biological levels during active social interactions. Behavioural synchrony between a parent and child manifests in the form of reciprocal gazes, expressions and vocalisations. Over time, these overt reciprocal behaviours become entrained into underlying patterns of synchrony in the brain that is unique to the dyad [12]–[15]. Biobehavioural synchrony in the parent-child dyad has been observed in the PFC. In one study, [16] revealed that synchrony was evident in the middle frontal gyrus (MFG) and superior frontal gyrus (SFG) of mother-child dyads when they played a game of Jenga together. Likewise, synchrony has been shown to occur in the dorsolateral PFC [15], the bilateral PFC and the temporo-parietal areas [13] of mother-child dyads during cooperative tasks. These findings suggest that biobehavioural synchrony that is unique to the parent-child relationship emerges in the PFC of dyads when they engage in coordinated interactions.

Despite extensive research on biobehavioural synchrony of parent-child dyads during active social interactions, few studies have investigated inter-subject synchronisation in dyads during everyday passive joint pursuits. Given the ubiquity with which parents and their children engage in co-viewing of narrative shows together in this digital age [17], it is imperative to understand whether such activities are associated with unique intersubject synchronisation in dyads, in which case such synchronisation could reflect partners’ attunement to each other in parent-child relationships. Previously, we have employed a co-viewing paradigm on mother-child dyads which showed that maternal parenting stress [18] and maternal anxious attachment [19] impairs intersubject synchronisation. Dyads in which mothers reported greater parenting stress showed diminished synchronisation in the medial left PFC whereas dyads with mothers who possess greater maternal attachment anxiety showed less synchronisation in the medial right PFC. The medial PFC is implicated in mentalisation processes which enables individuals to understand the mental state of others [20]. The findings from these previous synchronisation studies direct our attention to the medial region of the PFC as an area of the brain that is especially liable to display variations in inter-subject synchronisation during a co-viewing task.

Employing the same co-viewing paradigm as [18], the present study sought to examine whether father-child dyads would exhibit unique inter-subject synchronisation when co-viewing narrative animation scenes together. Hyperscanning functional Near-infrared Spectroscopy (fNIRS) would record the PFC activity of father-child pairs in tandem during the task. We embarked on this study with one central hypothesis. Given that the father-child relationship is an enduring form of early human attachment, we expected father-child dyads to exhibit unique inter-subject synchronisation, postulated to emerge in the medial region of the PFC, which would significantly differ from the synchronisation of two arbitrary adult-child participants.

## METHOD

### Participants

29 father-child dyads (11 girls, 18 boys) were recruited for this study through online platforms such as Facebook groups and forums (Mean age of fathers □ = □ 38.1years, ± 3.67 years; Mean age of children □ = □ 42.2 months, ± 5.25 months). Table 1 reports the sociodemographic information of fathers and their children. To be eligible for this study, fathers had to be at least 21 years old with a child aged between 36 and 48 months at recruitment. Both father and child were screened for any cognitive deficits as well as visual and hearing impairments so as to ensure that they were able to respond to the empirical tasks. Before the study commenced, informed consent was obtained from all participants (fathers signed on behalf of their child). Remuneration was provided at the end of the study to compensate for their gracious participation. All procedures were abided by relevant guidelines and were approved by the Institutional Review Board of Nanyang Technological University (IRB NTU-IRB-2018-06-016). All data are available at this URL: https://doi.org/10.21979/N9/PFHB88

**Table 1.** Sociodemographic information of fathers and children.

|  |  |  |
| --- | --- | --- |
| *Age of Fathers* | Mean age of 38.1 years (SD = 3.67) |  |
| *Age of Children* | Mean age of 42.2 months (SD = 5.25) |  |
| *Educational Level of Fathers* | 8 | Graduate/ Professional degrees |
|  | 12 | Standard college/ Bachelor's degrees |
|  | 2 | Partial college |
|  | 7 | Pre-college |
| *Employment* | 26 | Full-time |
|  | 1 | Part-time |
|  | 2 | Self-employed |
| *Household Income* | 9 | \$1,000 - \$4,999 |
| | 8 | \$5,000 - \$8,999 |
| | 5 | \$9,000 - \$12,999 |
| | 4 | \$13,000 - \$19,999 |
| | 3 | > \$20,000 |

### Experimental procedure

Father and child pairs were invited to the laboratory where they watched three video clips while their PFC activities were being recorded with functional Near-infrared Spectroscopy (fNIRS) in hyperscanning mode. The child sat on the father’s lap throughout the duration of the study (Fig. 1). NIRS caps of suitable sizes were fitted on the head of both the father and child. Optodes were adjusted and the signal quality was calibrated on the laptop prior to commencing with the study. As the devices were being set up, the child was distracted by a short 1-min video clip from the movie ‘Moana’ which was screened on a second laptop. This second laptop was placed approximately 40cm from the dyad and was also used to present the three video stimuli.

**Figure 1.**
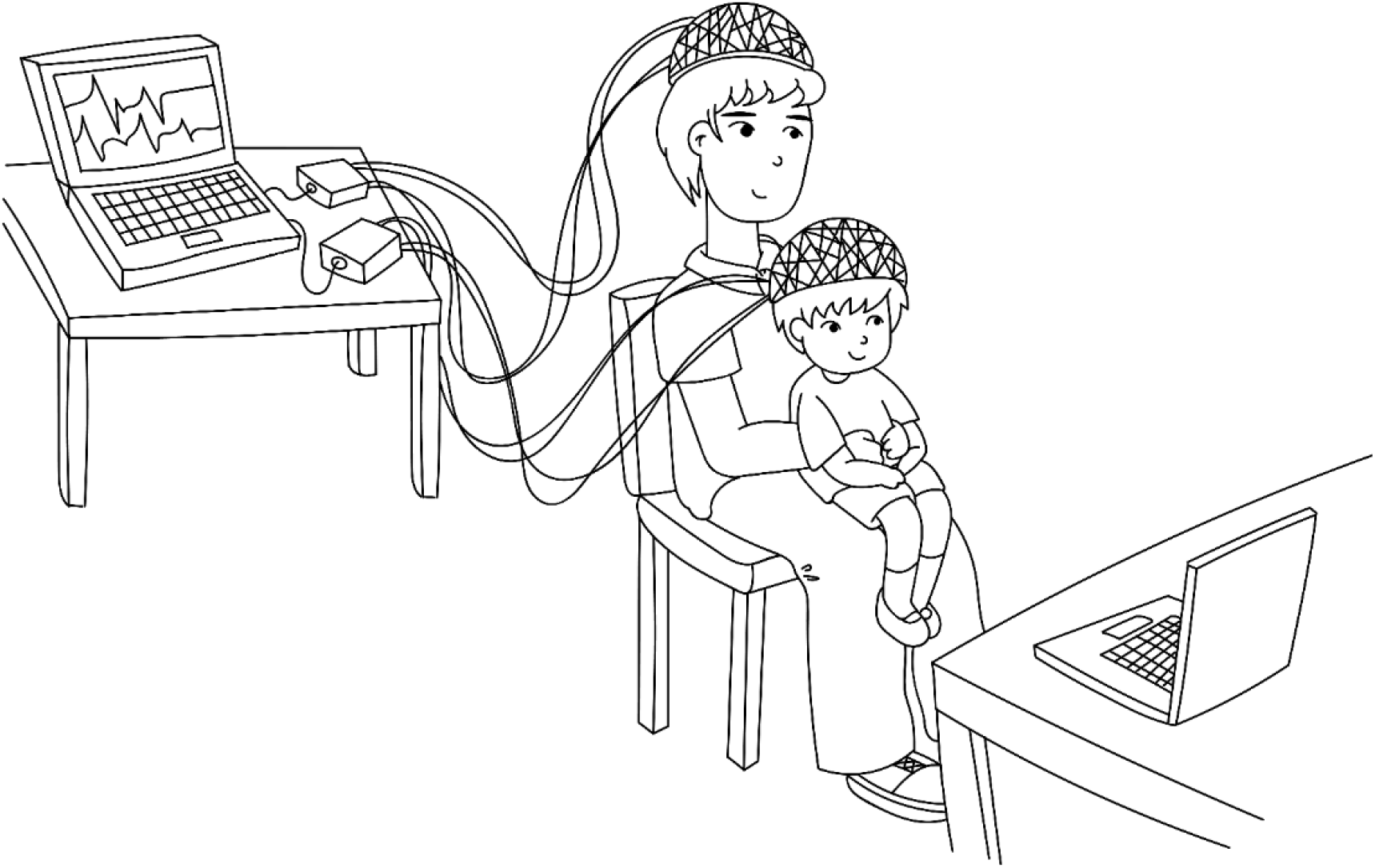
Schematic diagram and experimental set-up and seating arrangement of father and child. Figure illustrated by Farouq Azizan.

When the session was concluded, children rated how positive each of the video clips was on a visual analogue scale (VAS) which was administered via a Samsung Galaxy tablet. Fathers and children reported how frequently they watched these animation shows together. Finally, fathers were debriefed and remuneration was provided to the participants.

### Video stimuli

The video stimuli used in this study are the same as that from our previous study on mother-child dyads [18]. Father-child pairs were instructed to watch three 1-min excerpts of “Brave”, “Peppa Pig” and “The Incredibles”. To generalise the task of watching animation shows together, these clips were specifically chosen due to their different video complexity, audio fundamentals and audio intensity. The values of these parameters are reported in Table 2. Video complexity of each clip was analysed using Python and the FFmpeg software (v. 3.4.4) at a rate of 12 frames per second (FPS). To evaluate audio-related information, the video files were first converted into audio files using FFmpeg. Then, Praat software (v. 6.0.46) was used to measure audio intensity and audio fundamentals of the clips.

**Table 2.** Table reporting the video complexity, audio fundamentals, audio intensity, and positivity ratings of the three video clips.

| Video Stimulus | Video Complexity<br>(bytes) | Audio Fundamentals<br>(Hz) | Audio Intensity<br>(W/m<sup>2</sup>) |
| --- | --- | --- | --- |
| Brave | 658119.31 | 245.26 | 59.89 |
| Peppa Pig | 468369.81 | 218.37 | 62.50 |
| The Incredibles | 423005.66 | 271.64 | 56.30 |

The clips were edited to ensure that they were of similar volume and brightness. Prior to the onset of the first video clip, a 5-sec fixation cross was added. Between subsequent video clips, a 10-sec fixation cross was included (Fig. 2). The videos were presented in a pseudo-random order, where six different sequences of clips were first created before dyads were randomly assigned to one of the six sequences. The videos were screened on a 15-inch Acer Laptop in a dimly lit room. The brightness on the laptop was set to 165nits whereas the volume was at 44dB.

**Figure 2.**
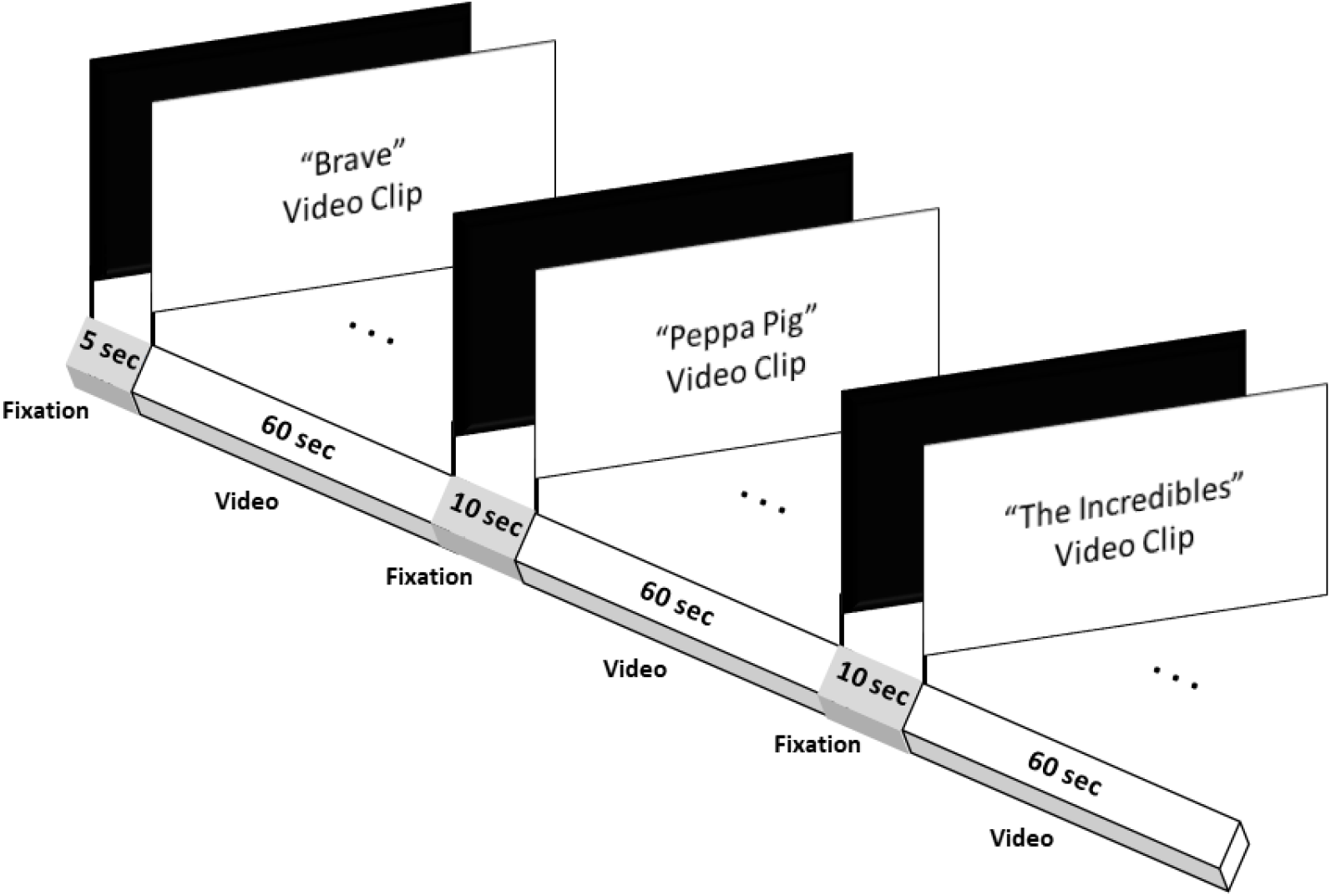
Schematic flow of stimuli presentation. The order in which the three video clips were screened to participants was pseudo-randomised.

### Functional near-infrared spectroscopy (fNIRS)

fNIRS was used to measure changes in concentration of oxygenated haemoglobin which served as a proxy of brain activation [21]. An fNIRS neuroimaging system (NIRSport, NIRx Medical Technologies LLC) with a 20-channel prefrontal cortex montage (NIRStar v.205 software), consisting of 8 sources and 7 detectors, was used in this study. LED sources of 760□nm and 850 nm wavelengths were applied at a scan rate of 7.81□Hz.

NIRSLab (nirsLAB v.2017.06) was used to preprocess the collected data. The preprocessing stream began with the setting of onset markers of the three video stimuli. Following that, motion correction of discontinuities and spike artefacts were conducted. Channels were then inspected for background noise, where those with Gain > 8 and CV > 7.5 were rejected and omitted from downstream preprocessing steps. To remove physiological and slow signals, a 0.01 Hz to 0.2 Hz bandpass filter was applied. The resulting signals were further inspected for undetected artifacts by two independent coders. Finally, the preprocessed optical signals were converted into changes in oxygenated (HbO) and deoxygenated (HbR) haemodynamic signals using the modified Beer-Lambert Law.

### Children’s Ratings of Video Stimuli

Upon the completion of fNIRS data acquisition, a Samsung Galaxy tablet was presented to child participants. To assess the positivity of the video stimuli, each child participant was asked to watch the three video clips again and select a facial expression that reflects the positivity of each clip from the visual analogue scale (VAS). The VAS presented a neutral face in the middle of a graphic slider. As the slider was moved one step to the left, the facial expression became more negative, and when the slider was moved two steps to the left from the centre position, the most negative facial expression was depicted. Conversely, when the slider was moved one step to the right, the facial expression became more positive, and moving the slider two steps to the right showed the most positive facial expression. The VAS was administered to the child with the help of the child’s father, where the father was asked the following question: “If you move the slider, you will notice that the mood of the face will change. By adjusting the slider, please ask your child to indicate whether he/she feels good or bad after watching the video.” The father then asks the child to click on the face which most reflects the emotional valence of the video. The facial expressions were later coded on a 5-point scale, where 1 denoted the most negative facial expression and 5 denoted the most positive facial expression.

To assess the familiarity of father-child dyads towards each video clip, fathers and their children were asked the following question: “How often do you watch this show?”. The dyads could choose from 4 options which were “Not at all”, “Not very often”, “Often” and “Very Often”. These responses were then coded on a 4-point scale, where 1 indicated no familiarity (i.e. “Not at all”) and 4 indicated extreme familiarity (i.e. “Very often”). Only data from 28 out of the 29 participants were recorded as one child participant refused to respond to the VAS.

### Cluster Signals

Similar to the previous study on mother-child dyads [18], PFC activity was analysed according to four clusters: frontal left, frontal right, medial left and medial right clusters (Fig. 3). For each subject, we derived the signals associated with the brain activity of each cluster, by computing the average of the normalised signals of the channels composing each cluster. The cluster brain activity signal is computed if at least 3 channel signals with good quality (each participant) are available. Therefore, the actual number of dyads used for the analysis is different for each video and cluster.

**Figure 3.**
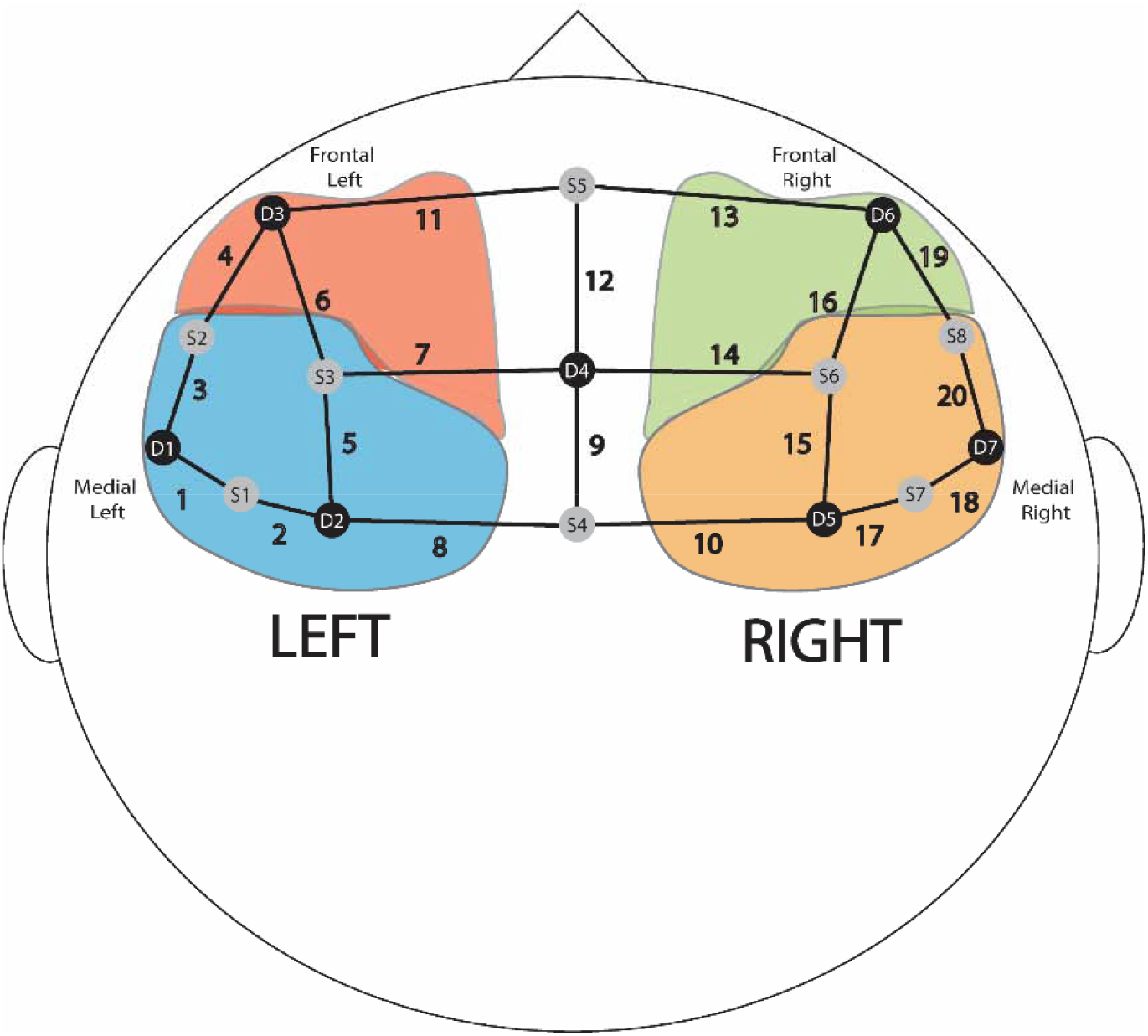
Schematic diagram of the four clusters of the prefrontal cortex: frontal left, frontal right, medial left and medial right. *Note: Grey dots represent sources, black dots represent detectors, and black lines represent channels. The number next to each line refers to the channel number*.

### Beta-Coefficient Analyses

Using NIRSLab (nirsLAB v.2017.06), a within-subject general linear model (GLM) was first performed so as to extract beta-coefficients for each of the three video stimuli, from each child and parent. The same GLM settings as [18] were used. Similarly, pre-whitening was omitted and a haemodynamic response function (HRF) was specified. A Discrete Cosine Transformation (DCT) temporal parameter with a 128 sec high-pass period cut-off and a Gaussian Full Width at Half Maximum (FWHM) 4 model were applied to the GLM. Beta-coefficients from each of the 20 channels were extracted from all participants. Within each participant, beta-coefficients were aggregated across the three video stimuli.

To account for different brain activation patterns in groups of fathers and that of children, analysis of covariance (ANCOVA) was first performed on each cluster, for fathers and children, separately. For each ANCOVA model, aggregated beta values were fitted as the dependent variable and video stimuli was included as the independent variable. Since mentalisation when viewing narrative scenes is influenced by age [22], [23], the ages of father and child were included as covariates. Children’s individual ratings of positivity and familiarity of the three videos were also incorporated as covariates in the model (i.e. Beta_Fathers_ ~ Video + Age_Fathers_ + Age_Children_ + Positivity Rating + Familiarity Rating; Beta_Children_ ~ Video + Age_Fathers_ + Age_Children_ + Positivity Rating + Familiarity Rating).

### Inter-subject Synchronisation Analyses

Cross Correlation within a maximum delay of 2 seconds [24]–[26] was used to compute synchrony between the normalised cluster signals, for each video and cluster. Two types of synchrony were computed:

- Control synchrony: between signals of all father-child dyads except the true dyads. This type of synchrony represents the effect of the stimulus (i.e. the effect arising from viewing the same narrative scenes);
- True synchrony: between the signals of the father and child of the true dyads. This type of synchrony represents both the effects of the stimulus and copresence (i.e. the effect arising from a true father and child being in each other’s presence).

Inter-subject synchronisation was preliminarily analysed using ANCOVA analyses as an attempt to account for the effects of covariates which emerged to be significant in the preliminary beta-coefficient analyses (i.e. Synchrony ~ Dyad (Control vs True Dyads) + Cluster + Dyad * Father’s Age + Dyad * Child’s Age + Dyad * Positivity).

Since the main research question of this study was to investigate the effect of copresence, a Mann-Whitney test was subsequently conducted to compare between True and Control synchrony in each cluster and in response to each video stimulus, without other modulating variables. This test would demonstrate that the copresence of a true father and child pair has a significant effect in increasing the inter-subject synchronisation of father-child brain activity. P-values were corrected for multiple hypotheses (False Discovery Rate correction Benjamini-Hochberg method).

## Results

### Children’s Ratings of Video Stimuli

Table 3 reports the means and standard deviations of children’s ratings of positivity for each video and the frequency of viewing these videos at home prior to the study.

**Table 3.** Table reporting the descriptive statistics of children’s positivity ratings of videos and frequency of viewing videos.*Note: Peppa refers to the scene from “Peppa Pig”and Incredibles refers to the scene from “The Incredibles”*.

| Ratings of Videos |  |  |  |  |
| --- | --- | --- | --- | --- |
|  | Positivity of Videos |  | Frequency of Viewing Videos |  |
|  | Mean | SD | Mean | SD |
| Brave | 4.464 | 0.962 | 1.357 | 0.559 |
| Peppa | 4.357 | 1.062 | 2.607 | 1.100 |
| Incredibles | 3.143 | 1.533 | 1.214 | 0.499 |

A dependent 2-group Wilcoxon signed rank test (one-sided) with continuity correction was then conducted to determine whether there are significant differences in ratings of positivity across video stimuli, and of the frequency of viewing these scenes at home. Wilcoxon signed rank test shows that the scenes from both “Peppa Pig” and “Brave” are rated to be significantly more positive than “The Incredibles” (Table 3A). This result was expected as the two scenes from “Peppa Pig” and “Brave” depicted positive family interactions whereas the scene from “The Incredibles” depicted an emotionally arousing family conflict.

**Table 3.**
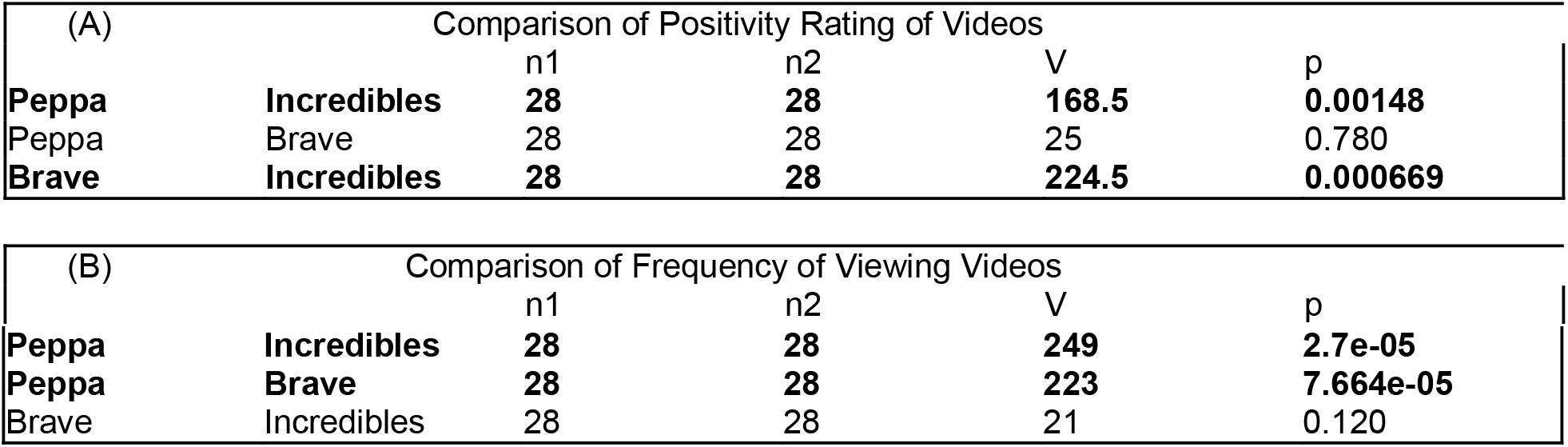
(A) Table reporting the statistical result from dependent 2-group Wilcoxon signed rank test (one-sided) comparing children’s ratings of positivity of videos. (B) Table reporting the statistical result from dependent 2-group Wilcoxon signed rank test (one-sided) comparing children’s ratings of frequency of viewing videos. *Note: Peppa refers to the scene from “Peppa Pig” and Incredibles refers to the scene from “The Incredibles”. Line in bold denotes significant comparisons after p-value correction*.

The test also revealed that scenes from “Peppa Pig” are viewed significantly more frequently compared to both “The Incredibles” and “Brave” (Table 3B). This result was also expected as “Peppa Pig” is a children’s television series which are possibly screened on the television more often whereas “The Incredibles” and “Brave” are both children’s movies.

### Beta-coefficient Results

ANCOVA analyses on fathers revealed a significant effect of child’s age and father’s age in the frontal right cluster, and a significant effect of child’s age in the medial left cluster (Table 4). Post-hoc Spearman’s correlation analyses later demonstrated that older fathers displayed greater activation in the frontal right cluster (p = 0.002, r_s_ = 0.54). Fathers with older children exhibited more activation in the frontal right cluster (p = 0.033, r_s_ = 0.397) and medial left cluster (p = 0.021, r_s_ = 0.428).

**Table 4.**
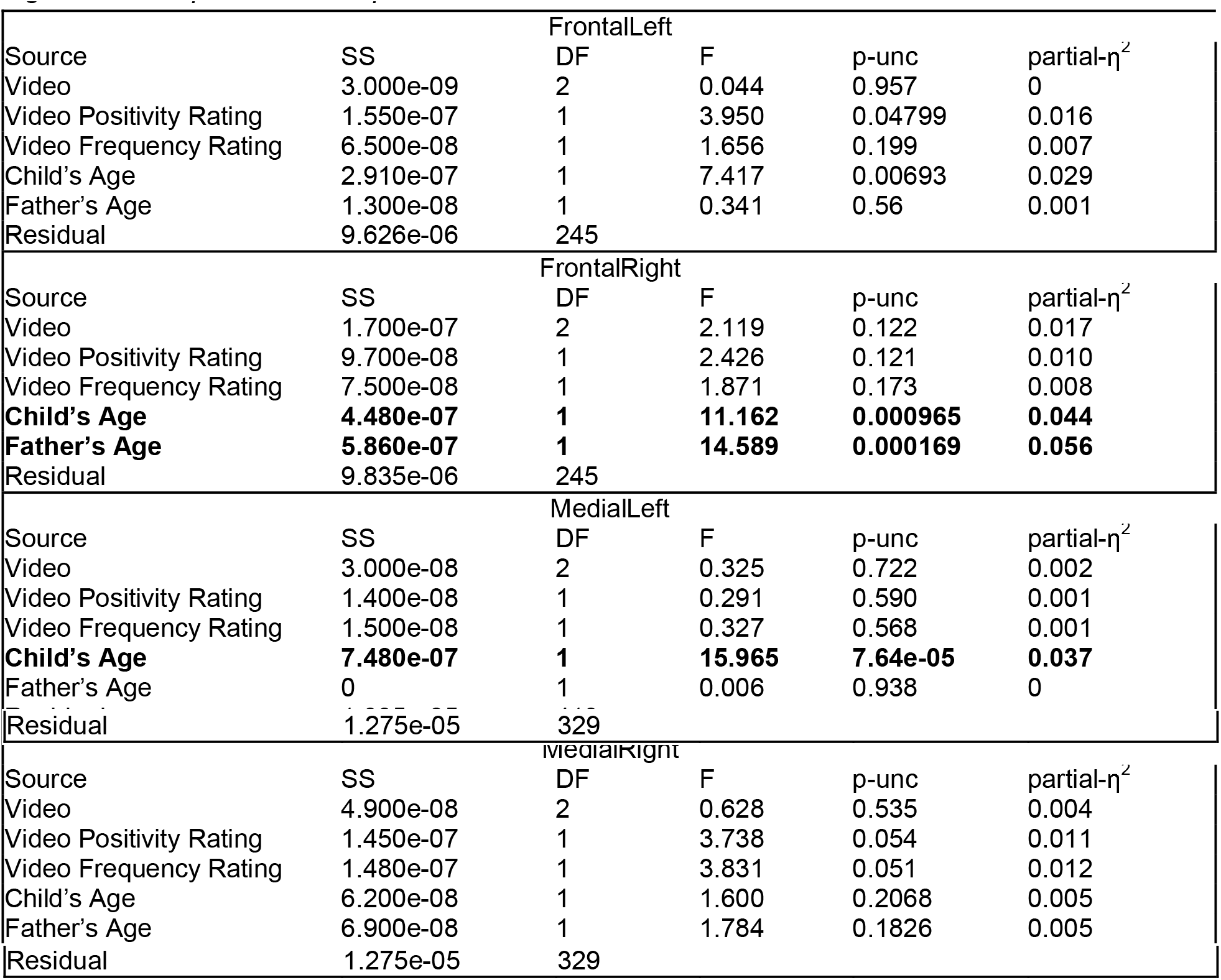
Table reporting the results of ANCOVA analyses on fathers. *Note: Line in bold denotes significant comparisons after p-value correction*.

| FrontalLeft |  |  |  |  |  |
| --- | --- | --- | --- | --- | --- |
| Source | SS | DF | F | p-unc | partial-$\eta^2$ |
| Video | 3.000e-09 | 2 | 0.044 | 0.957 | 0 |
| Video Positivity Rating | 1.550e-07 | 1 | 3.950 | 0.04799 | 0.016 |
| Video Frequency Rating | 6.500e-08 | 1 | 1.656 | 0.199 | 0.007 |
| Child's Age | 2.910e-07 | 1 | 7.417 | 0.00693 | 0.029 |
| Father's Age | 1.300e-08 | 1 | 0.341 | 0.56 | 0.001 |
| Residual | 9.626e-06 | 245 |  |  |  |
| FrontalRight |  |  |  |  |  |
| Source | SS | DF | F | p-unc | partial-$\eta^2$ |
| Video | 1.700e-07 | 2 | 2.119 | 0.122 | 0.017 |
| Video Positivity Rating | 9.700e-08 | 1 | 2.426 | 0.121 | 0.010 |
| Video Frequency Rating | 7.500e-08 | 1 | 1.871 | 0.173 | 0.008 |
| **Child's Age** | **4.480e-07** | **1** | **11.162** | **0.000965** | **0.044** |
| **Father's Age** | **5.860e-07** | **1** | **14.589** | **0.000169** | **0.056** |
| Residual | 9.835e-06 | 245 |  |  |  |
| MedialLeft |  |  |  |  |  |
| Source | SS | DF | F | p-unc | partial-$\eta^2$ |
| Video | 3.000e-08 | 2 | 0.325 | 0.722 | 0.002 |
| Video Positivity Rating | 1.400e-08 | 1 | 0.291 | 0.590 | 0.001 |
| Video Frequency Rating | 1.500e-08 | 1 | 0.327 | 0.568 | 0.001 |
| **Child's Age** | **7.480e-07** | **1** | **15.965** | **7.64e-05** | **0.037** |
| Father's Age | 0 | 1 | 0.006 | 0.938 | 0 |
| Residual | 1.935e-05 | 413 |  |  |  |
| MedialRight |  |  |  |  |  |
| Source | SS | DF | F | p-unc | partial-$\eta^2$ |
| Video | 4.900e-08 | 2 | 0.628 | 0.535 | 0.004 |
| Video Positivity Rating | 1.450e-07 | 1 | 3.738 | 0.054 | 0.011 |
| Video Frequency Rating | 1.480e-07 | 1 | 3.831 | 0.051 | 0.012 |
| Child's Age | 6.200e-08 | 1 | 1.600 | 0.2068 | 0.005 |
| Father's Age | 6.900e-08 | 1 | 1.784 | 0.1826 | 0.005 |

|  |  |  |
| --- | --- | --- |
| Residual | 1.275e-05 | 329 |

ANCOVA analyses on children showed a significant effect of video positivity rating in the medial right cluster (Table 5). Subsequent post-hoc Spearman’s correlation analysis revealed that a higher video positivity rating was associated with greater activation in the child’s medial right cluster (p = 0.016, r_s_ = 0.316).

**Table 5.** Table reporting the results of ANCOVA analyses on children. *Note: Line in bold denotes significant comparisons after p-value correction*.

| FrontalLeft |  |  |  |  |  |
| --- | --- | --- | --- | --- | --- |
| Source | SS | DF | F | p-unc | partial-$\eta^2$ |
| Video | 2.580e-07 | 2 | 2.078 | 0.127 | 0.017 |
| Video Positivity Rating | 7.500e-08 | 1 | 1.211 | 0.272 | 0.005 |
| Video Frequency Rating | 2.630e-07 | 1 | 4.229 | 0.041 | 0.017 |
| Child's Age | 6.000e-09 | 1 | 0.1 | 0.752 | 0 |
| Father's Age | 6.000e-08 | 1 | 0.962 | 0.328 | 0.004 |
| Residual | 1.523e-05 | 245 |  |  |  |
| FrontalRight |  |  |  |  |  |
| Source | SS | DF | F | p-unc | partial-$\eta^2$ |
| Video | 5.870e-07 | 2 | 4.527 | 0.012 | 0.036 |
| Video Positivity Rating | 9.900e-08 | 1 | 1.521 | 0.219 | 0.006 |
| Video Frequency Rating | 2.900e-08 | 1 | 0.446 | 0.505 | 0.002 |
| Child's Age | 4.600e-08 | 1 | 0.702 | 0.403 | 0.003 |
| Father's Age | 1.600e-07 | 1 | 2.467 | 0.118 | 0.010 |
| Residual | 1.588e-05 | 245 |  |  |  |
| MedialLeft |  |  |  |  |  |
| Source | SS | DF | F | p-unc | partial-$\eta^2$ |
| Video | 3.770e-07 | 2 | 2.817 | 0.061 | 0.013 |
| Video Positivity Rating | 2.940e-07 | 1 | 4.388 | 0.037 | 0.011 |
| Video Frequency Rating | 2.980e-07 | 1 | 4.447 | 0.036 | 0.011 |
| Child's Age | 6.800e-08 | 1 | 1.023 | 0.312 | 0.002 |
| Father's Age | 2.200e-08 | 1 | 0.333 | 0.564 | 0.001 |
| Residual | 2.763e-05 | 413 |  |  |  |
| MedialRight |  |  |  |  |  |
| Source | SS | DF | F | p-unc | partial-$\eta^2$ |
| Video | 5.180e-07 | 2 | 4.937 | 0.008 | 0.029 |
| **Video Positivity Rating** | **4.680e-07** | **1** | **8.920** | **0.003** | **0.026** |
| Video Frequency Rating | 2.600e-08 | 1 | 0.499 | 0.481 | 0.002 |
| Child's Age | 3.270e-07 | 1 | 6.231 | 0.013 | 0.019 |
| Father's Age | 7.100e-08 | 1 | 1.359 | 0.244 | 0.004 |
| Residual |  | 329 |  |  |  |

### Inter-subject Synchronisation Results

#### (i) Preliminary ANCOVA Analyses

Since child’s age, father’s age and video positivity rating were found to be significant from the beta analyses, these variables were included as covariates in the ANCOVA analysis of inter-subject synchrony. The ANCOVA analysis revealed a significant effect of Dyad (True vs Control) (Table 6). A post-hoc one-tailed Mann-Whitney U test later revealed that True dyads showed significantly more synchrony than Control dyads (U = 411660.0, p = 0.000503).

**Table 6.** Table reporting the statistical result from ANCOVA analysis of inter-subject synchrony. *Note: Line in bold denotes significant comparisons after p-value correction*.

| Source | SS | DF | F | p-unc | partial-$\eta^2$ |
| --- | --- | --- | --- | --- | --- |
| **Dyad (True vs Control)** | **0.706** | **1** | **10.727** | **0.001** | **0.003** |
| Cluster | 0.348 | 3 | 1.762 | 0.152 | 0.001 |
| **Father's Age** | **1.323** | **1** | **20.107** | **<0.001** | **0.005** |
| Child's Age | 0.327 | 1 | 4.969 | 0.026 | 0.001 |
| Video Positivity Rating | 0.010 | 1 | 0.149 | 0.700 | <0.001 |
| Dyad : Father's Age | 0.401 | 1 | 6.1 | 0.014 | 0.002 |
| Dyad : Child's Age | 0.003 | 1 | 0.038 | 0.845 | <0.001 |
| Dyad : Video Positivity Rating | 0.437 | 1 | 6.641 | 0.010 | 0.002 |
| Residual | 241.273 | 3668 |  |  |  |

The ANCOVA analysis also showed a significant main effect of father’s age. Subsequent Spearman’s correlation test between synchrony (both True and Control dyads) and father’s age demonstrated that the older the father, the less synchrony was observed in father-child pairs (r_s_ = −0.063, p = 0.000147).

#### (ii) Mann-Whitney Test

The Mann-Whitney test indicated that the for “The Incredibles” scene, the synchrony value in the medial left cluster for true dyads (Mean synchrony = 0.30, SD = 0.24) is significantly greater than that of control dyads (Mean synchrony = 0.10, SD = 0.25): U=1174, p=0.013, which means that the copresence of true father and child has a significant effect in modulating intersubject synchronisation towards the emotionally arousing “The Incredibles” scene (Table 7).

**Table 7.** Table reporting the statistical results from Mann-Whitney analysis which compared the synchrony value of True and Control dyads in each cluster and in response to each video stimulus. *Note: Peppa refers to the scene from “Peppa Pig” and Incredibles refers to the scene from “The Incredibles”. Line in bold denotes significant comparisons after p-value correction*.

| video | cluster | N | True dyads |  | Control dyads |  | U | p-uncorr | p-corr |
| --- | --- | --- | --- | --- | --- | --- | --- | --- | --- |
|  |  |  | Mean | SD | Mean | SD |  |  |  |
| Brave | FrontalLeft | 18 | 0.11 | 0.24 | 0.07 | 0.25 | 1492.0 | 0.282 | 0.484 |
| Brave | FrontalRight | 20 | 0.12 | 0.19 | 0.12 | 0.26 | 1908.0 | 0.488 | 0.559 |
| Brave | MedialLeft | 15 | 0.13 | 0.22 | 0.12 | 0.24 | 798.0 | 0.468 | 0.559 |
| Brave | MedialRight | 17 | 0.09 | 0.17 | 0.10 | 0.20 | 1151.0 | 0.513 | 0.559 |
| Peppa | FrontalLeft | 18 | 0.17 | 0.25 | 0.10 | 0.24 | 1588.0 | 0.145 | 0.347 |
| Peppa | FrontalRight | 20 | 0.21 | 0.26 | 0.17 | 0.25 | 2007.0 | 0.340 | 0.510 |
| Peppa | MedialLeft | 15 | 0.22 | 0.22 | 0.17 | 0.27 | 881.0 | 0.230 | 0.461 |
| Peppa | MedialRight | 17 | 0.10 | 0.31 | 0.13 | 0.27 | 1127.0 | 0.568 | 0.568 |
| Incredibles | FrontalLeft | 17 | 0.20 | 0.24 | 0.12 | 0.24 | 1342.0 | 0.141 | 0.347 |
| Incredibles | FrontalRight | 20 | 0.21 | 0.21 | 0.11 | 0.26 | 2321.0 | 0.052 | 0.208 |
| **Incredibles** | **MedialLeft** | **15** | **0.30** | **0.24** | **0.10** | **0.25** | **1174.0** | **0.001** | **0.013** |
| Incredibles | MedialRight | 17 | 0.26 | 0.25 | 0.15 | 0.28 | 1457.0 | 0.041 | 0.208 |

## Discussion

The present study aimed to investigate whether father-child dyads would exhibit unique inter-subject synchronisation during co-viewing of narrative scenes. Only one hypothesis was postulated which was that true father-child dyads would display significantly greater synchrony than control dyads in the medial PFC. This hypothesis was partially supported since preliminary analyses that considered demographic and stimuli covariates, such as the age of participants and the perceived positivity and familiarity of the videos, evinced that true dyads showed greater inter-subject synchronisation than control dyads in the PFC as a whole, rather than the medial region only. However, when comparing the synchrony values of true and control dyads only, true dyads indeed displayed greater synchrony than control dyads in the medial left PFC cluster, but this effect was only observed in response to the emotionally arousing conflict scene from “The Incredibles”.

Our preliminary attempt at investigating inter-subject synchronisation, while controlling for demographic and stimuli covariates, demonstrated that father-child dyads show unique inter-subject synchronisation during co-viewing of movie stimuli. This finding implies that being in a father-child attachment relationship confers greater coordination in processing narrative scenes compared to arbitrary adult-child pairs. [9] proposed that inter-subject synchronisation occurs across arbitrary individuals to some degree due to the activation of a common script which enables the similar interpretation of narrative scenes. However, displaying significantly greater synchronisation than arbitrary pairs of individuals suggests that the processing of narrative scenes in father-child dyads were specifically modulated by each other’s presence. Nestled within an attachment relationship, father-child dyads are familiar with each other’s social experiences and interactional environments [10], [11]. Hence, one postulation is that co-viewing could have activated a script which is unique to the dyad that facilitates greater synchronisation in their joint processing of narrative scenes. For instance, the scene from “The Incredibles” portrayed characters who were engaged in a heated argument at home. Dyadic partners could have interpreted the scene similarly depending on how common, or uncommon, such a conflict scenario is in the family. Likewise, the scene from “Brave”, which depicted family members hugging each other elatedly, could be processed by father-child pairs using a unique script that is shaped by whether such overt displays of affection commonly occur in the family.

The subsequent analysis that focused solely on the difference between true and control dyads showed that the effect of dyadic presence on inter-subject synchronisation emerged in the left medial PFC region in response to the conflict scene of “The Incredibles”. Viewing any movie stimulus would require the recruitment of mentalisation processes in the medial PFC so as to understand the mental and emotional states of characters in the narrative [27], [28]. However, since greater synchronisation in true compared to control dyads was observed in “The Incredibles” scene only, the emotional intensity of the scene being attended to could be a factor in eliciting dyad-specific synchronisation. [2] similarly showed that greater inter-subject synchronisation was observed across arbitrary participants during emotionally arousing segments of the movie. [5] later postulated this event to be due to the vicarious activations of the brain when observing how another individual, bearing a similar physical human form, undergoes the same intense emotional experiences as one’s self. The inter-subject synchronisation observed by [2] during emotionally intense scenes could be further amplified in father-child dyads in this study due to the attunement that dyadic partners have towards each other’s emotional states that likely supersedes arbitrary vicarious activations in the brain. As father-child pairs are cognisant of each other’s behavioural repertoire, dyads could have been more attuned to subtle changes in each other’s social cues, such as an increase in tension of partners’ sitting posture, that convey emotional arousal. These subtle changes in the physical state of partners’ bodies could have then elicited similarities in brain processing which was reflected as unique inter-subject synchronisation.

Both synchrony and beta-coefficient analyses corroborate in revealing a main effect of father’s age. Dyads with older fathers exhibited diminished synchrony, and older fathers were also observed to display greater activation in the frontal right cluster compared to their younger counterparts. Although conclusive interpretations from these observations cannot yet be drawn, we present the postulation that age confers parents with a greater sense of security in their parenting role [30]–[32] which enabled older fathers to process narrative scenes in the presence of their child without feeling the need to excessively attune to their child’s emotional state. Another study on father-mother synchrony by [29] similarly found synchrony to be diminished in older couples in response to salient human vocalisations, including child-related cries and laughter. The authors suggested that older parents are likely more confident in their parenting roles due to their extensive life experiences, which could render them more discerning as to when they need to attune to their partners’ emotional state. These results generally suggest that parenting brain mechanisms are potentially modulated by parents’ age and stage of life.

Analysis of children’s beta-coefficients revealed that an enhanced activation in the medial right cluster of the PFC was associated with children’s ratings of video positivity. Child participants rated clips from “Brave” and “Peppa Pig” to be significantly more positive than the conflict scene from “The Incredibles”. Taken together, these findings seem to suggest that greater mentalisation activity in the medial right cluster was elicited in response to positive narrative scenes. Children’s proclivity for mentalising positive scenes could be derived from their greater exposure to positive social environments. In early childhood, preschool-aged children tend to be exposed to positive social experiences more so than negative ones [33]–[35], and this prior experience in positive situations could have facilitated mentalisation of positive narrative scenes. In comparison, preschool-aged children’s lack of negative social experiences could have posed a challenge for them to relate to the conflict scene from “The Incredibles” which was thus reflected as less mentalisation activity in the medial right PFC during this stimulus.

Several limitations ought to be assessed in this study. First, this paradigm used only one scene from each video stimuli which did not allow investigation of emotional valence within the same narrative. For example, since only a negative conflict scene from the unfamiliar movie “The Incredibles” was screened to the children, we were not able to disentangle the effects of a positive scene casted within the same unfamiliar narrative. Future studies should screen several positive and negative scenes in both familiar and unfamiliar narrative contexts. Second, the area of the brain that was investigated was only restricted to the prefrontal cortex.

Inter-subject synchronisation could have spanned across other cortical and subcortical regions which belong to specific networks that are implicated in mentalisation processes. Some examples of brain areas involved in mentalisation include cortical structures such as the temporal cortex and subcortical areas like the amygdala [36]-[38]. Future studies may adopt more extensive brain montages when conducting inter-subject synchronisation studies. Third, this study did not include a manipulation to determine the effects of physical contact between the father and child. Having the child on the father’s lap could have incurred a significant effect of touch on synchronous brain responses. Future studies should also examine synchrony in dyads when they are seated side-by-side to ensure that touch is controlled in the experimental design.

## Conclusion

In this study, we sought to answer the question of whether father-child dyads would display unique inter-subject synchronisation during a typical co-viewing activity. Our preliminary synchrony analyses showed that father-child dyads exhibit unique inter-subject synchronisation, even upon controlling for demographic and stimuli variables. When comparing explicitly between true and control dyads, findings demonstrated that dyad-specific synchronisation was observed in the medial left region of the prefrontal cortex when father-child pairs viewed an emotionally arousing scene together. This finding provides significant insight to our understanding of how an early and enduring attachment, such as that of a parent-child relationship, elicits a temporal coordination of mentalisation events that possibly reflects partners’ attunement to each other. Since synchrony was observed to diminish in older fathers, future research should also investigate the role of parental age and stage of life in modulating synchrony in parent-child relationships.

## Acknowledgements

This work was supported by the NAP Start-up Grant M4081597 (G.E.) from Nanyang Technological University Singapore as well as the Ministry of Education Tier-1 Grant RG55/18 (NS) 2018-T1-001-172 (G.E.), and the Singapore Children’s Society (A.A.). The founding agencies had no role in the conceptualization, design, data collection, analysis, decision to publish, or preparation of the manuscript. We would like to extend our appreciation to Justin Randall Durnford, Jan Paolo Macapinlac Balagtas, Siti Syazana Binti Abdul Halim, Anais Ang, Jezebel Chin Syuen Chong, Wan Ting Wong, Michelle Neoh and Lim Meng Yu for their assistance in the project, and Farouq Azizan for his illustration.

## Contributions

A.A. and G.E. conceptualized the study; A.A. collected and pre-processed the data. A.A. and A.B. performed all analyses. A.A. wrote the original draft; G.E., A.A., and A.B. reviewed and edited the submitted version of the article.

